# Convolutional neural network models of neuronal responses in macaque V1 reveal limited non-linear processing

**DOI:** 10.1101/2023.08.26.554952

**Authors:** Hui-Yuan Miao, Frank Tong

## Abstract

Computational models of the primary visual cortex (V1) have suggested that V1 neurons behave like Gabor filters followed by simple non-linearities. However, recent work employing convolutional neural network (CNN) models has suggested that V1 relies on far more non-linear computations than previously thought. Specifically, unit responses in an intermediate layer of VGG-19 were found to best predict macaque V1 responses to thousands of natural and synthetic images. Here, we evaluated the hypothesis that the poor performance of lower-layer units in VGG-19 might be attributable to their small receptive field size rather than to their lack of complexity *per se*. We compared VGG-19 with AlexNet, which has much larger receptive fields in its lower layers. Whereas the best-performing layer of VGG-19 occurred after seven non-linear steps, the first convolutional layer of AlexNet best predicted V1 responses. Although VGG-19’s predictive accuracy was somewhat better than standard AlexNet, we found that a modified version of AlexNet could match VGG-19’s performance after only a few non-linear computations. Control analyses revealed that decreasing the size of the input images caused the best-performing layer of VGG-19 to shift to a lower layer, consistent with the hypothesis that the relationship between image size and receptive field size can strongly affect model performance. We conducted additional analyses using a Gabor pyramid model to test for non-linear contributions of normalization and contrast saturation. Overall, our findings suggest that the feedforward responses of V1 neurons can be well explained by assuming only a few non-linear processing stages.

## Introduction

The primary visual cortex is arguably the best understood cortical area and has served as a critical testbed for developing neurocomputational models of visual processing. Since the pioneering work of Hubel and Wiesel (1962), researchers have sought to characterize and understand the tuning properties of V1 simple cells and complex cells (Mechler & Ringach, 2002; Priebe, 2016; Priebe et al., 2004; Ringach et al., 2002). Simple cells respond to preferred orientations in a position-specific manner, whereas complex cells are believed to pool information from multiple simple cells that share a common orientation preference. Early conceptual models of V1 neuronal tuning (Hubel & Wiesel, 1962) helped set the foundation for subsequent computational models. Jones and Palmer (1987) modeled simple cell responses by using 2D Gabor filters to account for a neuron’s orientation, spatial frequency, and phase tuning preferences. Likewise, complex cells can be modeled by squaring and summing the outputs from a quadrature pair of linear filters to obtain phase-invariant responses to stimuli (Adelson & Bergen, 1985). Subsequent work has shown that the response properties of simple and complex cells vary along a continuum (Mechler & Ringach, 2002; Ringach et al., 2002), and that the outputs from multiple linear filters can be combined to obtain better predictions of V1 responses (Rust et al., 2005; Vintch et al., 2015). For example, a top-performing convolutional subunit model consisted of multiple spatially shifted copies of a single linear filter, followed by non-linear rectification, weighted pooling of these subunit responses, and a final non-linearity applied to the pooled response (Vintch et al., 2015). A key property of these models is that they require only a few non-linear processing stages to account for V1 neuronal responses.

By contrast, a recent study by Cadena et al. (2019) has suggested that V1 neurons perform far more non-linear computations than previously expected. The authors used the layer-wise unit activity of a deep convolutional neural network (CNN), VGG-19 (Simonyan & Zisserman, 2015), to predict neuronal responses in macaque V1 to thousands of natural and synthetic images (see **Figure 1**). VGG-19 performs convolutional filtering followed by non-linear rectification in each convolutional layer, and max-pooling operations every few layers, to learn effective representations for object classification. VGG-19 was found to outperform traditional V1 models, and more surprisingly, the best performance occurred not in the lower layers of the CNN, but rather, in an intermediate layer (conv3_1) following five convolutional operations and two max-pooling operations. These findings led the authors to conclude that a large number of non-linear operations are needed to fully capture the response properties of V1 neurons. If true, this would imply that our understanding of the neural computations performed within area V1 was widely off the mark and that a major reconceptualization is required.

**Figure 1.**
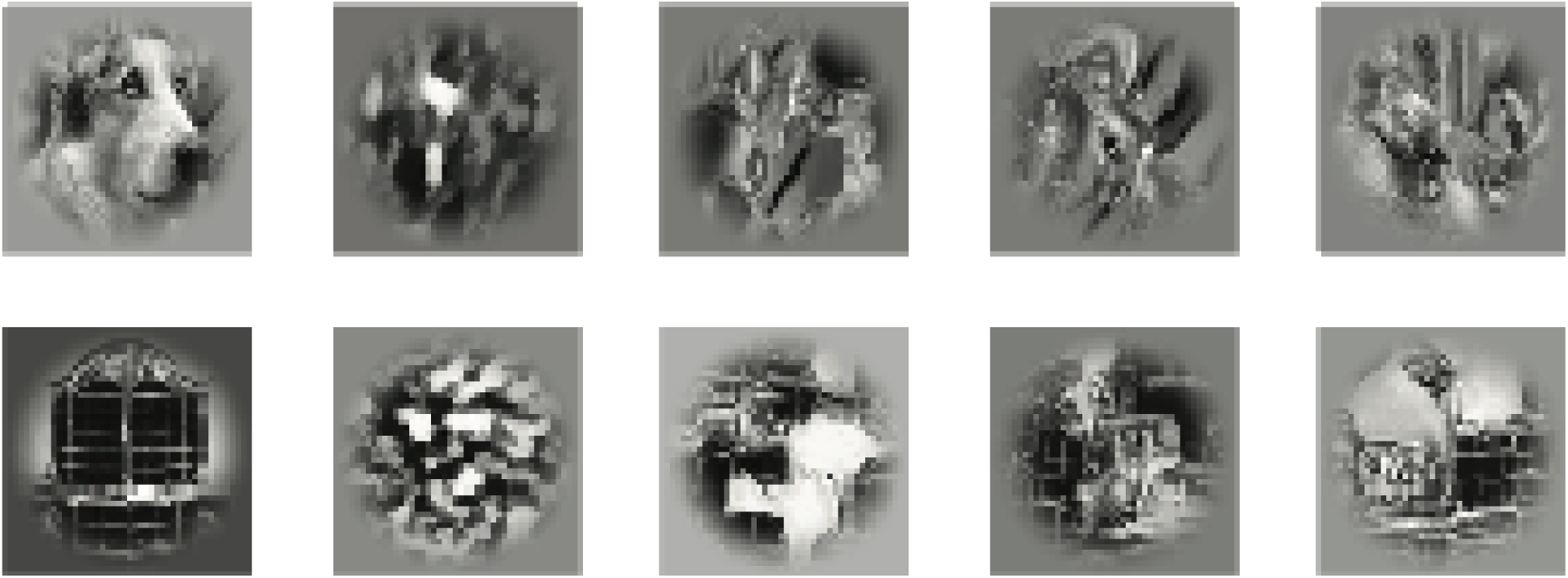
Examples of the stimuli used in this study. Leftmost column, examples of two natural images. Columns 2-5, synthetic images derived from the natural images to evoke similar responses in the lower layers of VGG-19, with increasing correspondence across multiple layers shown from left to right.

However, given the complexity of deep CNN models, it may not always be obvious as to why one CNN layer performs better at predicting neural responses in a particular visual area when compared to another layer (Guclu & van Gerven, 2015; Khaligh-Razavi & Kriegeskorte, 2014; Schrimpf et al., 2018; Yamins et al., 2014). The visual representations that are learned by the convolutional layers of a CNN are constrained not only by the number of non-linear operations that occur by a given stage of processing, but also by hyperparameters such as filter size and stride that determine the effective receptive field size of the units in each layer. In particular, VGG-19 relies on small convolutional filters (3×3 pixels) that are sampled by the subsequent layers with a stride of 1, leading to very small receptive fields in the lower layers (see **Table 1**). We hypothesized that the mismatch between the image size used to predict V1 responses (40×40 pixels) and the receptive field size of units in the lower layers of VGG-19 may have caused systematic biases in model performance to favor of higher layers with larger receptive fields, as fewer CNN units would then be needed to predict or account for a V1 neuron’s response properties.

**Table 1.** The architecture of standard VGG-19. The table shows the output dimensionality of each CNN layer and the associated kernel size, stride length and receptive field size in pixel units.

| Layer type | Output size | Kernel size | Stride | RF size |
| --- | --- | --- | --- | --- |
| Convolution (conv1_1) | 224×224×64 | 3 | 1 | 3 |
| Convolution (conv1_2) | 224×224×64 | 3 | 1 | 5 |
| Max pool (pool_1) | 112×112×64 | 2 | 2 | 6 |
| Convolution (conv2_1) | 112×112×128 | 3 | 1 | 10 |
| Convolution (conv2_2) | 112×112×128 | 3 | 1 | 14 |
| Max pool (pool_2) | 56×56×128 | 3 | 2 | 16 |
| Convolution (conv3_1) | 56×56×256 | 3 | 1 | 24 |
| Convolution (conv3_2) | 56×56×256 | 3 | 1 | 32 |
| Convolution (conv3_3) | 56×56×256 | 3 | 1 | 40 |
| Convolution (conv3_4) | 56×56×256 | 3 | 1 | 48 |
| Max pool (pool_3) | 28×28×256 | 2 | 2 | 52 |
| Convolution (conv4_1) | 28×28×512 | 3 | 1 | 68 |
| Convolution (conv4_2) | 28×28×512 | 3 | 1 | 84 |
| Convolution (conv4_3) | 28×28×512 | 3 | 1 | 100 |
| Convolution (conv4_4) | 28×28×512 | 3 | 1 | 116 |
| Max pool (pool_4) | 14×14×512 | 2 | 2 | 124 |
| Convolution (conv5_1) | 14×14×512 | 3 | 1 | 156 |
| Convolution (conv5_2) | 14×14×512 | 3 | 1 | 188 |
| Convolution (conv5_3) | 14×14×512 | 3 | 1 | 220 |
| Convolution (conv5_4) | 14×14×512 | 3 | 1 | 252 |
| Max pool (pool_5) | 7×7×512 | 2 | 2 | 268 |
| Fully connected layers for classification |  |  |  |  |

To address these concerns, we compared the neural predictivity of VGG-19 with AlexNet (Krizhevsky et al., 2012), which has much larger receptive fields in its first few layers. Surprisingly, we found that the best predictions of V1 responses arose from the first convolutional layer of a standard pre-trained version of AlexNet, in sharp contrast from VGG-19. Since the predictive accuracy of pre-trained AlexNet was slightly lower than the best-performing layer of VGG-19, we trained and tested a modified version of AlexNet that was designed to provide more feature maps in the lower layers to match the large number of feature representations available in conv3_1 of VGG-19. These analyses revealed that the first pooling layer of modified AlexNet performed just as well as VGG-19 at predicting V1 responses on an image-by-image basis. In a series of control analyses, we further show that the best-performing layer of VGG-19 was highly dependent on the size of the input images used to predict V1 responses.

Given that pre-trained AlexNet exhibited the highest neural predictivity in the first convolutional layer, which involves only a single stage of filtering and non-linear rectification, we were motivated to compare such CNN performance with that of a simple V1 Gabor pyramid model with hand-engineered rather than learned filters. We further generated and tested multiple versions of the V1 Gabor pyramid model to test for multiple effects of divisive normalization, including cross-orientation inhibition and surround suppression (Carandini et al., 1997). These analyses revealed a small but statistically significant benefit of incorporating normalization into these simpler V1 models. Although the Gabor V1 models performed slightly but significantly worse than the best CNN models we tested (i.e., conv3_1 of VGG-19 and pool_1 of modified AlexNet), these computational models are simpler and might be argued to provide a more parsimonious account of V1 neuronal tuning.

Taken together, our findings demonstrate that while V1 processing is not strictly linear, only a small number of non-linear computations are needed to account for the stimulus-driven response properties of the primary visual cortex.

## Materials and Methods

### Neurophysiology Data Set

The V1 neuronal data and the visual stimuli used in the original study were obtained from a publicly available data repository posted by Cadena et al. (2019). Below, we provide a brief description of their experimental design and methods; further details can be found in the original paper.

Two awake-behaving monkeys were presented with a set of 7250 complex images while activity from multiple isolated V1 neurons was recorded using a 32-channel linear silicon probe. The monkeys were trained to fixate on a small dot presented in the center of the display throughout each trial. Each trial consisted of a randomized sequence of 29 images, each shown for 60 ms. The images were centered over the population receptive field of the recorded neurons. A total of 250 trials were required to present all 7250 images in a pseudorandomized order while activity from multiple isolated units was recorded. The authors recorded responses from a total of 307 neurons over 23 recording sessions. Recordings of suitable quality were collected from 166 neurons with two or four independent observations obtained for all 7250 images; these data were used for the main analyses.

The image set consisted of 1450 natural images selected from the ImageNet database (Russakovsky et al., 2015) and four different sets of synthetic textures (four synthetic textures for each individual natural image) that were designed to evoke activation patterns in different convolutional layers of VGG-19 that closely resembled those elicited by the original natural images (Gatys et al., 2015). The natural images were first transformed into 256 × 256 grayscale images, then used to generate the synthetic images. Then, for all the natural and synthetic images, a circular crop was applied (140-pixel diameter, corresponding to 2° of visual angle), followed by a contrast-attenuated windowing procedure so that the image gradually faded beyond a diameter of 1° of visual angle (see **Figure 1**).

### Image preprocessing

For our study, the following image preprocessing steps were performed prior to analysis by the CNNs. For the images shown to the monkeys, the central 80 × 80 pixel region of the 140 × 140 pixel-sized images were cropped and then rescaled to 40 × 40 pixels using bicubic interpolation to match the input size used by Cadena et al. (2019). Bicubic interpolation was also chosen for all other image resizing procedures in this project. All grayscale images were z-scored before being fed to the models. The processed images were converted to RGB format and zero padding was applied if required by the CNN architecture.

### Building V1 model with CNN features

We modeled the V1 data set using two different approaches. In our first pipeline, we evaluated how well CNNs could predict V1 neuronal responses using linear regression, which was implemented using custom-written code in MATLAB. For the second pipeline, we adopted Cadena et al. (2019)’s computational approach, which relied on TensorFlow and stochastic gradient descent to fit a generalized linear model that assumed a Poisson noise distribution. Both pipelines relied on L1 regularization (Tibshirani, 1996) to select a sparser set of predictive units, which substantially improved model performance. Both pipelines led to very similar results and comparable levels of predictive performance. Here, we report the results of the second approach to facilitate the comparison of findings across studies.

To model V1 neuronal responses, we first presented the entire set of 7250 images to the CNNs to calculate the layer-wise activation patterns in each CNN layer. We used 80% of the images for model training (training set), and the remaining 20% of the images served as the independent test set. Image assignment was arbitrary, with the only constraint that any original image and its four synthetic derivatives were assigned to a common set.

We extracted the layer-wise activations corresponding to the image inputs after the application of rectified linear unit (ReLU) processing, as well as the unit activity found within the pooling layers. The activations at each individual layer were used to construct a generalized linear model (GLM) for each neuron in the V1 dataset to model the responses of individual neurons to the images in the training set. Before building the GLM model, we first applied batch normalization to the responses from each layer, such that the activity patterns obtained from each feature map were adjusted to have zero mean and unit variance across all images within a batch (batch size, 256 images); this procedure ensured that L1 normalization would apply the same level of penalty to all inputs. After batch normalization, we passed the activation map of a single CNN layer, ***X***, through **Equation 1** for which ***W*** is the weight matrix and ***B*** is the bias term. To guarantee that the model predictions were non-negative and to maintain consistency with Cadena et al.’s approach, we applied **Equation 2** to the output of **Equation 1**. L1 regularization was applied to improve model performance and stability with a lambda of 0.03 (determined via cross-validation using training data only). Unlike Cadena et al. (2019), we chose not to apply smoothness or group sparsity as these regularization factors had a negligible impact on the goodness of fit of the models as was also noted by Cadena et al. (2019).

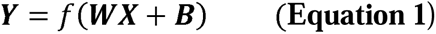

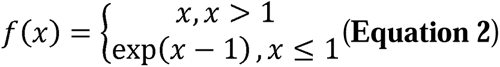

The models and regression analyses were implemented in Tensorflow (Abadi et al., 2016) following the approach of Cadena et al. (2019), and all model training was identical to Cadena unless otherwise stated. The model was optimized using the Adam optimization algorithm (Kingma & Ba, 2015) with a learning rate of 10^-4^ and a batch size of 256. An early stopping technique was used to determine when to cease model training. To implement the early stopping process, we used 80% of the images *within* the training set to tune the model parameter estimates, and the remaining portion of the training data to evaluate model performance every 100 training steps. If the performance did not improve for ten consecutive evaluations, the learning rate was reduced to 1/3 of its current value and this procedure was repeated three times.

### Model evaluation

After the regression weights were calculated using the training data, we evaluated the predictive accuracy of each CNN layer by calculating the Pearson correlation coefficient between the predicted and actual responses of individual neurons across all test images. Actual responses were based on the average number of spikes that were observed across different repetitions of a given test image for an isolated neuron. For model comparison, Fisher z-transformation was applied to the Pearson correlation scores obtained for each of the 166 neurons, after which we applied paired t-tests to determine which CNN models performed better at predicting neuronal responses.

### Noise ceiling

Given that each image was presented only a few times to each monkey (2 or 4 trials/image), it would be challenging to obtain reliable estimates of the lower and upper bounds of the noise ceiling. Nevertheless, we thought it would be informative to compute these estimates as they might provide some benchmark as to how well the best models are performing. We focused on analyzing the neurons with four measurements per image, since the neurons with only 2 observations per image would lead to an enormous gap between the lower and upper noise ceiling estimates. For the lower bound, we adopted a leave-one-trial-out approach and used the averaged response of the remaining 3 trials to predict the response observed for the left-out trial; this was performed on each of the 4 trials. This analysis revealed a lower bound of mean *r* = 0.3944. For the upper bound, we calculated the correlation between the mean response to an image across all 4 trials with the response observed on each individual trial. This analysis revealed an upper bound of mean *r* = 0.6811. We then compared the performance of our CNN models with the lower bound of the noise ceiling. Note that these correlation values are somewhat lower than the main text, since the CNN model is being used to predict responses to images on individual trials. We found that the predictive accuracy of the best layer of VGG-19 and modified AlexNet was slightly but significantly lower than the lower bound of the noise ceiling (VGG-19, *r* = 0.3791, *t*(114) = 2.45, p = 0.02; modified AlexNet: *r* = 0.3809, *t*(114) = 2.28, p = 0.02). Thus, while CNN-based models perform very well at accounting for these V1 responses, there may still be some room for further improvement.

### Comparisons between AlexNet and VGG-19

We first compared the performance of VGG-19, using the same model version as Cadena et al. (2019), with that of AlexNet by employing the pre-trained version available in PyTorch (Paszke et al., 2019). The layer that provided the best predictions of V1 neuronal responses was identified for each model, and performance accuracy (i.e., Pearson correlation) was then compared across the two models. The best-performing layer of VGG-19, conv3_1, consisted of 256 feature maps where each unit had an effective receptive field size of 24 pixels that corresponded to 0.69° of visual angle in the neurophysiological experiments. The large number of feature maps available at conv3_1 could potentially provide an undue advantage for regression-based models to fit the responses of individual V1 neurons. Thus, in addition to our evaluation of a pre-trained AlexNet model, we evaluated the performance of a modified AlexNet architecture that incorporated additional convolutional layers to support a larger number of feature representations in the lower layers of the CNN (see **Table 2**). We also made modest adjustments to the filter sizes of this model so that the receptive field size would increase more gradually after layer 1. We trained modified AlexNet on RGB images (224 × 224 × 3 px) using the 1000 categories of images from ImageNet (ILSVRC2012). Modified AlexNet was trained using stochastic gradient descent over a period of 100 epochs with an initial learning rate of 0.01 (decreased by a factor of 10 every 30 epochs), a batch size of 256, a weight decay of 0.0005, and a momentum of 0.9. Modified AlexNet reached a top-1 accuracy of 0.51. A similar training regime was used to train the modified VGG-19 models, with more training epochs used to train this deeper network. Modified VGG-19 models were trained using stochastic gradient descent over a period of 180 epochs with an initial learning rate of 0.01 (decreased by a factor of 10 every 60 epochs), a batch size of 256, a weight decay of 0.0001, and a momentum of 0.9. Version 1 of VGG-19 attained a top-1 accuracy of 0.54, and version 2 attained a top-1 accuracy of 0.57.

**Table 2.** Comparison between the architectures of standard AlexNet and modified AlexNet.

| Layer type | AlexNet |  |  |  | Modified AlexNet |  |  |  |
| --- | --- | --- | --- | --- | --- | --- | --- | --- |
|  | Output size | Kernel size | Stride | RF size | Output size | Kernel size | Stride | RF size |
| Convolution | 55×55×64 | 11 | 4 | 11 | 75×75×64 | 11 | 3 | 11 |
| Convolution | - | - | - | - | 75×75×256 | 5 | 1 | 23 |
| Max pool | 27×27×64 | 3 | 2 | 19 | 37×37×256 | 2 | 2 | 26 |
| Convolution | 27×27×192 | 5 | 1 | 51 | 37×37×512 | 3 | 1 | 38 |
| Convolution | - | - | - | - | 37×37×512 | 3 | 1 | 50 |
| Max pool | 13×13×192 | 3 | 2 | 67 | 18×18×512 | 2 | 2 | 56 |
| Convolution | 13×13×384 | 3 | 1 | 99 | 18×18×512 | 3 | 1 | 80 |
| Convolution | 13×13×256 | 3 | 1 | 131 | 18×18×512 | 3 | 1 | 104 |
| Convolution | 13×13×256 | 3 | 1 | 163 | 18×18×512 | 3 | 1 | 128 |
| Max pool | 6×6×256 | 3 | 2 | 195 | 9×9×512 | 2 | 2 | 140 |
| Fully connected layers for classification |  |  |  |  |  |  |  |  |

### Evaluation of a V1 Gabor pyramid model

We also evaluated the performance of a Gabor-based V1 model, both to serve as a baseline comparison with the more complex CNN models and to assess whether V1 responses to complex natural images show evidence of divisive normalization (Carandini & Heeger, 2012). The Gabor wavelet pyramid (Lee, 1996) consisted of Gabor filters that occurred at spatial frequencies of 1, 2, 4 or 8 cycles per field of view (40 × 40 px) with a specified bandwidth of 1 octave. At each spatial frequency *f*, the Gabor filters were positioned on a 2*f* × 2*f* grid. At each grid location, there was a set of Gabor filters with eight possible orientations (0°, 22.5°, 157.5°) and four spatial phases (0°, 90°, 180°, 270°). The filters were truncated by setting the proportion of the filters with a value less than 1% of the peak value to 0. All filters were normalized to have zero mean and unit length. We simulated simple cell responses by applying half-wave rectification (or ReLU) to the responses of the Gabor units for all four spatial phases. To simulate complex cell responses, the outputs from Gabor filters with phase values of 0° and 90° were squared, summed, and then square-rooted to calculate these responses. The responses of all units were rescaled by the (near-)maximum possible response that they might produce if they were presented with a square-wave grating at the optimal spatial frequency, orientation, and spatial phase.

In comparison to the basic Gabor pyramid model, we evaluated whether the incorporation of contrast saturation (or contrast response non-linearity) would lead to better prediction of V1 responses by applying an exponentiation function (see **Equation 3**).

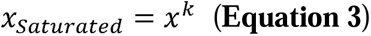

Given the response of a simulated unit, *x*, a contrast-saturated response can be generated by applying an exponent factor *k* using parameter values less than 1. We evaluated this contrast-saturated Gabor model’s performance using exponent values ranging from 0.1 to 1.0 (with 0.1 increments) and found that an exponent value of 0.6 led to the highest level of performance.

In addition, we simulated the effects of divisive normalization to see whether the performance of the Gabor-based V1 model could be further improved. Three types of divisive normalization were simulated (**Equation 4**): one with spatially restricted divisive normalization (mimicking cross-orientation inhibition), one with orientation-tuned normalization from the surround, and one with non-selective normalization from the surround.

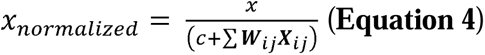

Given the response of a simulated unit, *x*, we could calculate a normalized response by incorporating the linearly weighted sum of the responses of that unit and its relevant neighbors in the denominator of **Equation 4**. Here, ***X****_ij_* represents the matrix of responses from all units in the normalization pool, ***W****_ij_* represents weight assigned to each of these responses, and *c* is a single parameter that controls the strength of divisive normalization. In all cases, normalization was performed separately on simple cell responses and complex cell responses.

For spatially restricted normalization, the normalization pool consisted of units with any orientation preference that shared the same preferences for spatial location and spatial frequency. We tested a range of values for parameter *c* ranging from 0.01 to 2.0 and found that a value of 0.25 led to the best model performance.

We evaluated two different models of surround suppression; both relied on a 2D Gaussian function to weight the responses of neighboring units in a spatially graded manner that declined as a function of distance. For the orientation-tuned surround suppression model, the normalization pool consisted of units that shared the same orientation and spatial frequency preferences. By contrast, the non-selective surround suppression model incorporated the responses of neighboring units with any orientation preference as long as they were tuned to the same spatial frequency. The spatial size of the normalization pool was scaled relative to the receptive field size of the simulated unit. Specifically, the standard deviation of the 2D Gaussian used to generate the linear weighting function (***W****_ij_*) was obtained by incorporating the standard deviation of the Gabor unit’s receptive field and then scaling that value by a multiplicative factor (α). We evaluated a range of values of α (from 0.1 to 1.5) to identify those that led to the best model performance (α = 0.75 for orientation-selective surround model, α = 0.75 for non-selective surround model). Likewise, we varied the parameter value for *c* (ranging from 0.05 to 1.0) to find those that led to the best performance for the orientation-tuned surround suppression model (*c* = 0.25) and the non-selective surround suppression model (*c* = 0.1).

Finally, we evaluated the impact of applying divisive normalization after first incorporating contrast saturation (using an exponent of 0.6) to determine whether the performance of the Gabor-based V1 model could be further improved. We used a similar approach as described above to identify parameter values for *α* and *c* that led to the best model performance. Because of the non-linear impact of contrast saturation on the simulated responses, we expanded the range of *c* (from 0.05 to 2.5) and the range of α (from 0.05 to 1.5) to identify the parameter settings that led to the best performance.

### Code accessibility

Following the publication of this article, the code used in the reported analyses will be made publicly available at https://github.com/Huiyuan-Miao/V1-Nonlinear.

## Results

### Evaluation of CNN models

We compared the layer-wise performance of VGG-19 and AlexNet in terms of their ability to predict V1 neuronal responses to a large set of complex grayscale images (see **Materials and Methods**). The full image set consisted of 1450 natural images and an additional 5800 synthetic images that were generated to mimic the low-level properties of the natural images; every image was presented twice to one monkey and four times to a second monkey.

We sought to determine whether AlexNet would necessarily require multiple non-linear processing steps to attain peak predictive performance, as was the case for VGG-19. The standard AlexNet architecture relies on much larger filters in layer 1 (11 × 11 pixels) and a stride value of 4 to sample from the input images; these factors lead to much larger receptive field sizes in the first few convolutional layers of AlexNet (**Table 2**) when compared to VGG-19 (**Table 1**).

We first replicated the main findings of Cadena et al. (2019) by confirming that the activation map of conv3_1 (the fifth convolutional layer) in VGG-19 provided the best predictions of V1 neuronal responses (**Figure 2A**). For simplicity of comparison, we adopted the same regression-based modeling approach of Cadena et al. by using unit activations in each layer as predictors of a V1 neuron’s response to a set of training images (80% of the data), with L1 regularization to estimate the weights for a sparse set of predictor units obtained from each layer. The trained model was then used to predict the neuron’s response to an independent set of test images (20% of the data).

**Figure 2.**
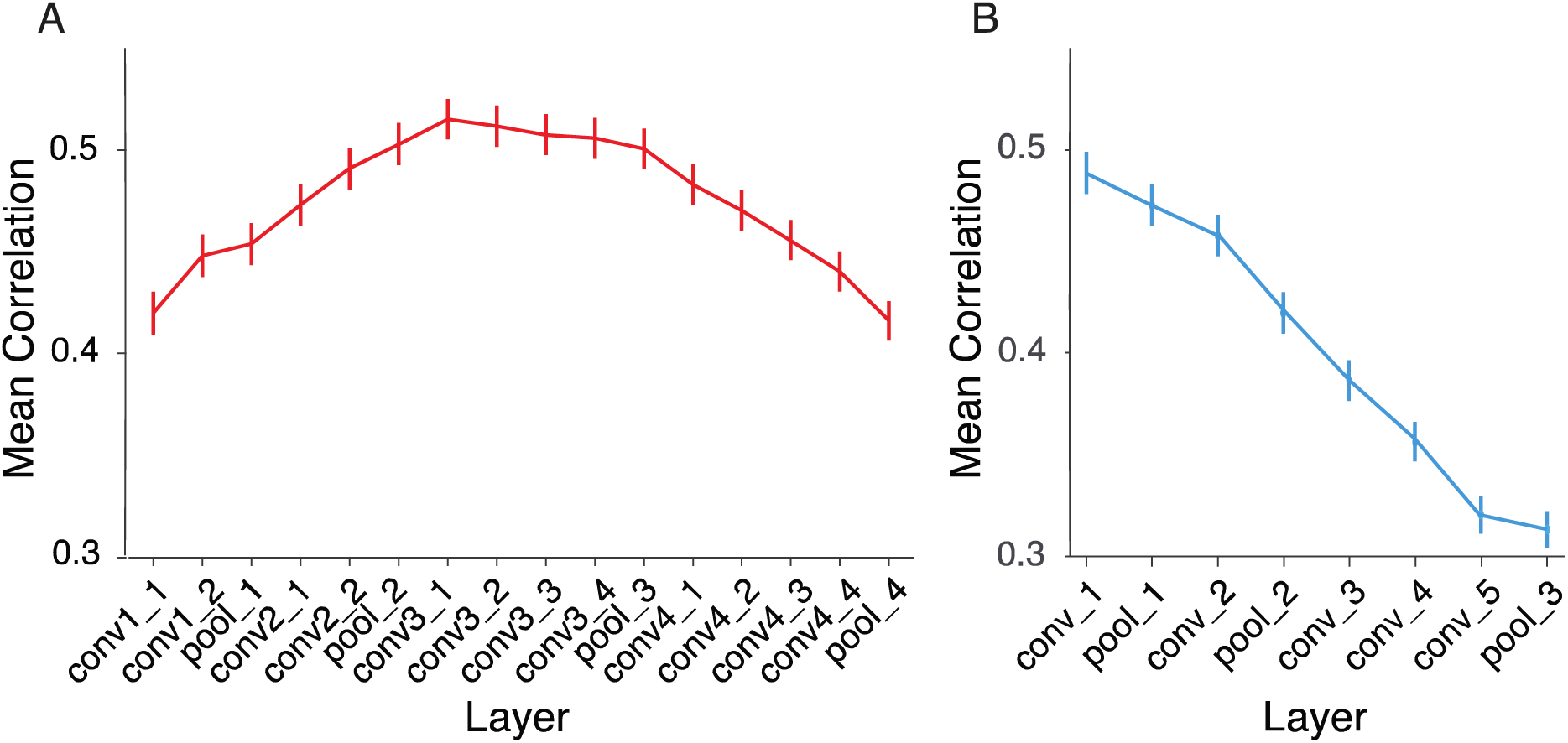
Layer-wise model predictions of V1 responses for standard versions of VGG-19 (A) and AlexNet (B). Plots show the mean correlation between the predicted and actual responses of 166 neurons, using the patterns of unit activity from individual CNN layers as regressors. Best predictive performance emerges at a much later processing stage for VGG-19 (conv3_1) when compared to AlexNet (conv_1). Error bars indicate ±1 standard error of the mean.

Of potential relevance, the receptive field size of the units in conv3_1 (24 × 24 pixels) spanned approximately half of the width of the input images (40 × 40 pixels) that were originally used to calculate CNN model responses, whereas prior stages of convolutional processing consisted of units with comparatively smaller receptive fields ranging from 3 to 16 pixels in width. It is worth noting that the input images used to generate VGG-19 model responses would have spanned only 1.14° of visual angle in the macaque V1 recording study, which recorded neuronal activity from parafoveal regions of V1 at eccentricities ranging from 1.4° to 3.0° of visual angle.

In sharp contrast with the layer-wise performance of VGG-19, we found that the best predictive performance of AlexNet emerged in the first convolutional layer (**Figure 2B**) with performance declining in a monotonic fashion for each processing stage thereafter. These findings indicate that AlexNet requires far fewer non-linear processing stages to obtain its best possible fit of V1 neuronal responses. However, it was still the case that the best-performing layer of VGG-19 showed significantly higher predictive accuracy than the best-performing layer of AlexNet (mean *r* of 0.5158 vs. 0.4887 respectively, averaged across 166 V1 neurons, *t*(165) = 10.09, *p* = 6.02×10^-19^). Thus, it remained possible that the many additional non-linear computations performed by VGG-19 were still required to attain this higher level of predictive accuracy.

We suspected that the poorer performance of the standard pre-trained version of AlexNet might be attributable to a few factors. One consideration was that the first convolutional layer of AlexNet had receptive fields that were considerably smaller than the input images used for model evaluation (about 1/4 in size, see **Table 2**), whereas the receptive fields for convolutional layer 2 (51 × 51 pixels) and all subsequent layers exceeded the size of the input images. We wondered whether the best possible predictive performance might be easier to attain if the CNN receptive fields are only modestly smaller than that of the input images, leading to a so-called *Goldilocks* zone. A second consideration was the fact that conv3_1 of VGG-19 has a very large number of feature channels (i.e., 256 channels) in comparison to conv1 of AlexNet (i.e., 64 channels), which might have conferred an advantage to VGG-19 by providing a larger number of predictors or basis functions to account for more subtle aspects of a V1 neuron’s receptive field structure.

We were therefore motivated to construct a modified architecture for AlexNet, which involved inserting an additional convolutional layer before the first pooling layer and increasing the number of feature channels in conv_2 and subsequent layers (see **Table 2** for a complete description of our modifications). This CNN model was then trained to classify the 1000 object categories from ImageNet (Russakovsky et al., 2015) until classification accuracy reached an asymptotic level of performance after 100 epochs (see **Materials and Methods** for details).

We performed the same analyses on this modified version of AlexNet and found that V1 predictivity improved across convolutional layers 1 and 2, peaking at the first max-pooling layer (r = 0.5167), after which performance steadily declined (**Figure 3A**). A direct comparison of the predictive accuracy of pool_1 of modified AlexNet with conv3_1 of VGG-19 (**Figure 3B)** indicated that the two CNN models did not reliably differ in their performance (*t*(165) = 0.680, *p* = 0.497). These findings rule out the possibility that V1 neuronal responses can only be adequately explained by considering a much more complex set of non-linear computations, such as those performed by the level of conv3_1 of VGG-19. Instead, our findings indicate that the number of non-linear steps needed to account for V1 responses is unlikely to be as large as that suggested by Cadena and colleagues (2019).

**Figure 3.**
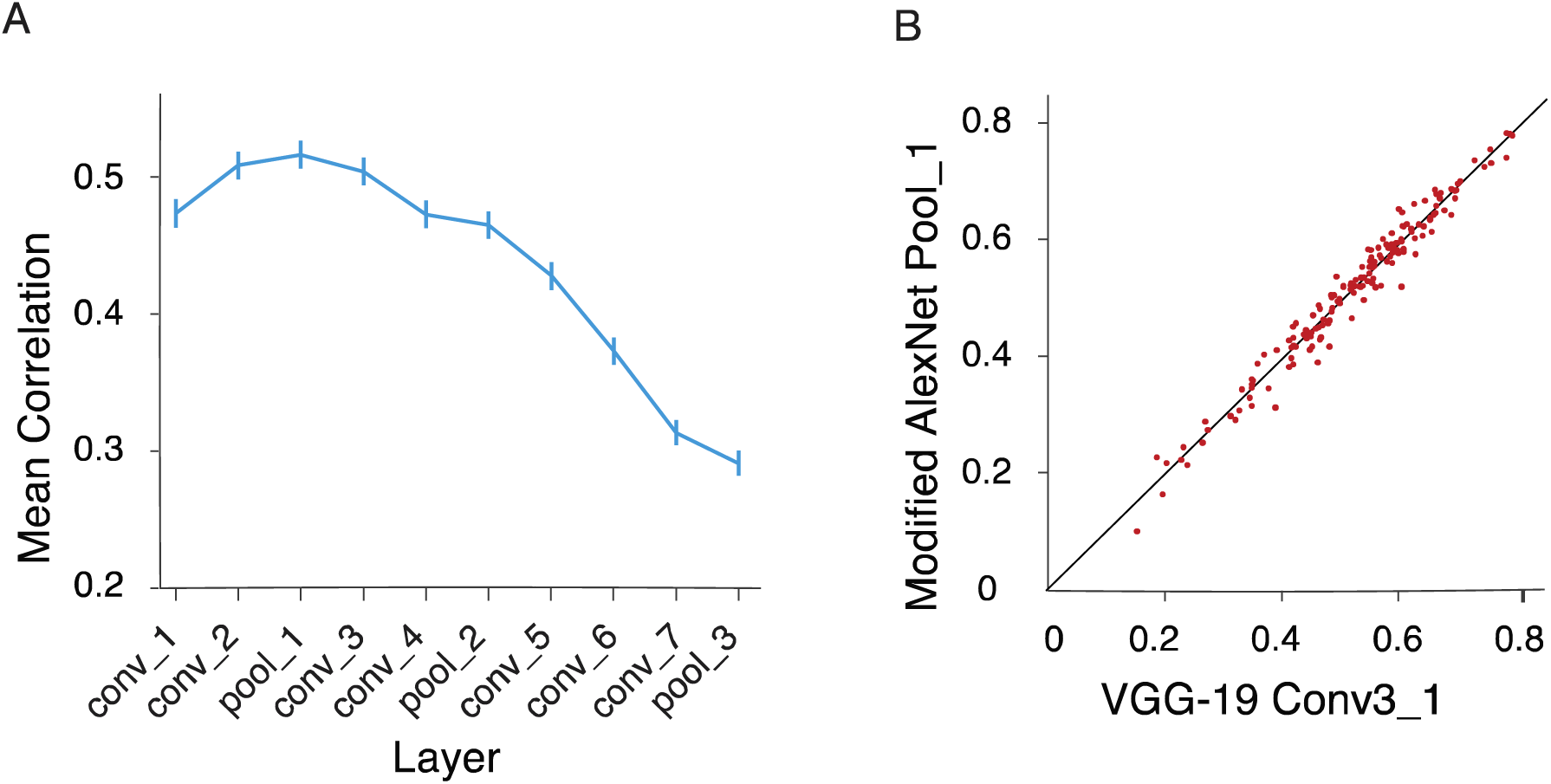
Model performance of modified AlexNet. A) Mean correlation between predicted and actual V1 neuronal responses is highest at the first pooling layer (pool_1) and thereafter decreases monotonically. B) Comparison of neural predictivity for the best-performing layer of VGG-19 and modified AlexNet. Each plot symbol indicates the prediction accuracy for a single V1 neuron. The performance of modified AlexNet’s pool_1 layer is comparable to that of VGG-19 conv3_1.

If the V1 predictivity of a CNN layer depends at least partly on the relationship between CNN receptive field size and the size of the input images used to generate the model’s responses, then one would expect that manipulations of input size should lead to systematic shifts in the layer that can yield the best performance. Specifically, smaller input images should lead to better performance in the lower or earlier layers of a CNN, whereas larger input images should lead to better performance in higher layers.

To test this hypothesis, we evaluated the performance of VGG-19 using input images that were scaled to be smaller (20 × 20 pixels) or larger (80 × 80 pixels) than the input size originally used for model evaluation (i.e., 40 × 40 pixels). As can be seen in **Figure 4**, this analysis confirmed that a smaller input size caused the best performance to shift to a lower layer of VGG-19 (i.e., pool_2 or the 6^th^ layer), whereas the larger input size biased performance to favor a much higher layer (i.e., conv4_1 or the 12^th^ layer). These findings demonstrate how non-neural factors, distinct from the complexity or number of computational operations used for modeling, can impact the ability of a CNN model to predict neuronal responses.

**Figure 4.**
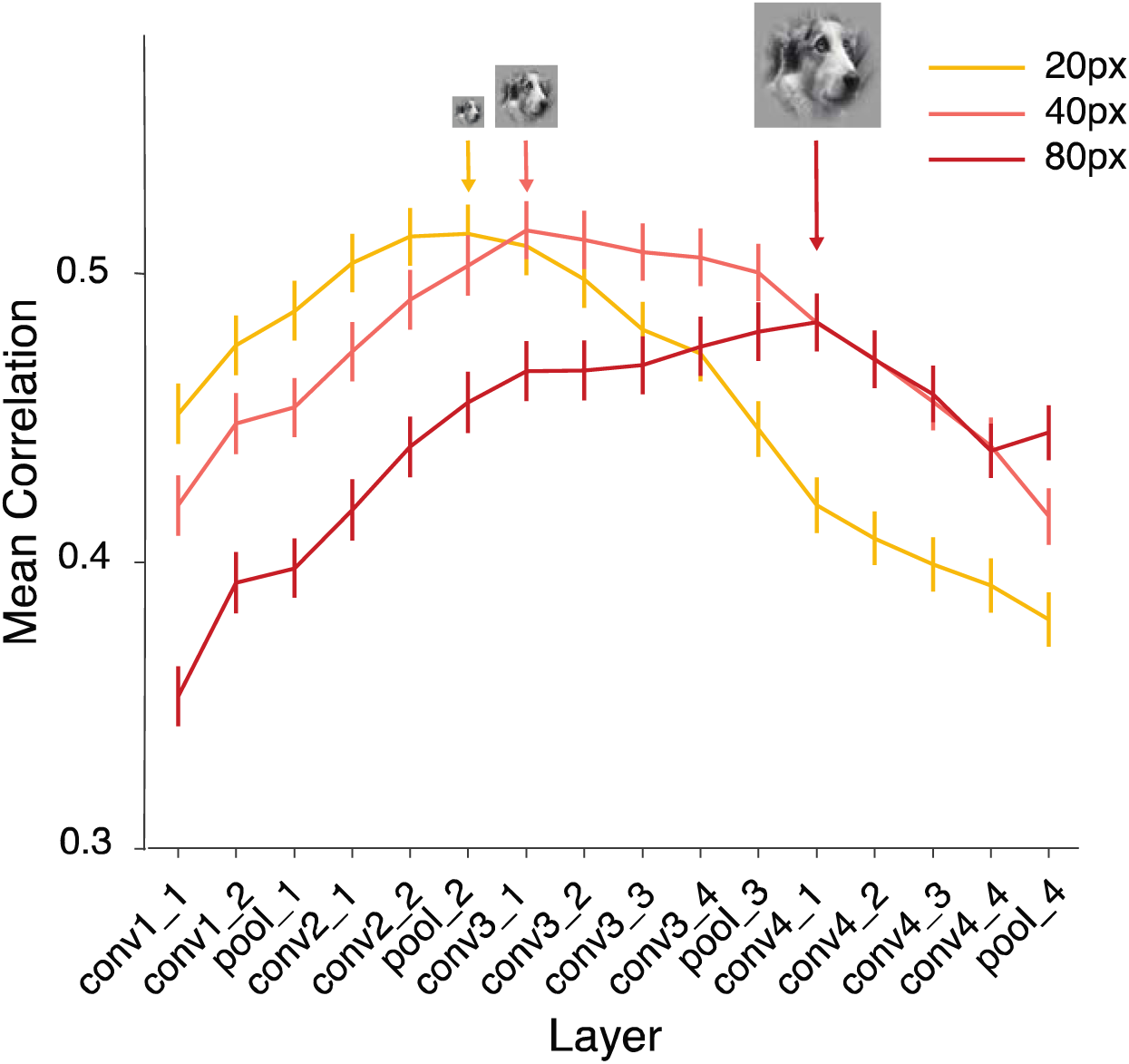
Model performance of VGG-19 when tested with different input image sizes. Images were scaled to pixel dimensions of 20 × 20, 40 × 40 (original analysis) or 80 × 80. An increase in input image size caused the best performance of VGG-19 to shift to higher layers.

As a final test of the idea that receptive field size can strongly affect which layer of a CNN can attain the best predictive accuracy, we constructed two modified versions of VGG-19 that had larger receptive fields in their lower layers. Specifically, these models sampled from convolutional layer 1 or layers 1-2 with increased stride length (see **Table 3**). After the models were trained on ImageNet object classification, their layer-wise performance was evaluated.

**Table 3.**
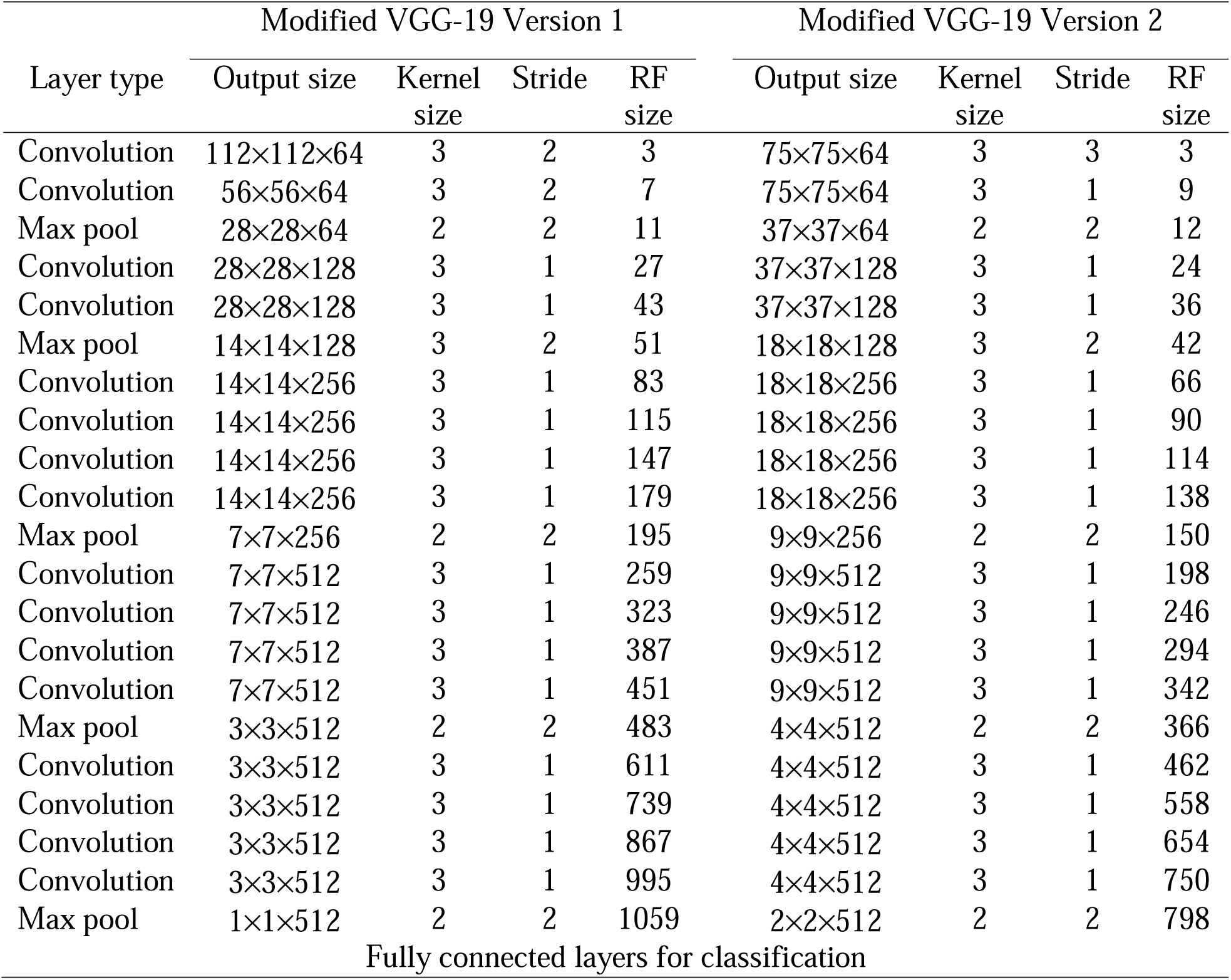
The architecture of the modified VGG-19 Version 1 and Version 2.

| Layer type | Modified VGG-19 Version 1 |  |  |  | Modified VGG-19 Version 2 |  |  |  |
| --- | --- | --- | --- | --- | --- | --- | --- | --- |
|  | Output size | Kernel size | Stride | RF size | Output size | Kernel size | Stride | RF size |
| Convolution | 112×112×64 | 3 | 2 | 3 | 75×75×64 | 3 | 3 | 3 |
| Convolution | 56×56×64 | 3 | 2 | 7 | 75×75×64 | 3 | 1 | 9 |
| Max pool | 28×28×64 | 2 | 2 | 11 | 37×37×64 | 2 | 2 | 12 |
| Convolution | 28×28×128 | 3 | 1 | 27 | 37×37×128 | 3 | 1 | 24 |
| Convolution | 28×28×128 | 3 | 1 | 43 | 37×37×128 | 3 | 1 | 36 |
| Max pool | 14×14×128 | 3 | 2 | 51 | 18×18×128 | 3 | 2 | 42 |
| Convolution | 14×14×256 | 3 | 1 | 83 | 18×18×256 | 3 | 1 | 66 |
| Convolution | 14×14×256 | 3 | 1 | 115 | 18×18×256 | 3 | 1 | 90 |
| Convolution | 14×14×256 | 3 | 1 | 147 | 18×18×256 | 3 | 1 | 114 |
| Convolution | 14×14×256 | 3 | 1 | 179 | 18×18×256 | 3 | 1 | 138 |
| Max pool | 7×7×256 | 2 | 2 | 195 | 9×9×256 | 2 | 2 | 150 |
| Convolution | 7×7×512 | 3 | 1 | 259 | 9×9×512 | 3 | 1 | 198 |
| Convolution | 7×7×512 | 3 | 1 | 323 | 9×9×512 | 3 | 1 | 246 |
| Convolution | 7×7×512 | 3 | 1 | 387 | 9×9×512 | 3 | 1 | 294 |
| Convolution | 7×7×512 | 3 | 1 | 451 | 9×9×512 | 3 | 1 | 342 |
| Max pool | 3×3×512 | 2 | 2 | 483 | 4×4×512 | 2 | 2 | 366 |
| Convolution | 3×3×512 | 3 | 1 | 611 | 4×4×512 | 3 | 1 | 462 |
| Convolution | 3×3×512 | 3 | 1 | 739 | 4×4×512 | 3 | 1 | 558 |
| Convolution | 3×3×512 | 3 | 1 | 867 | 4×4×512 | 3 | 1 | 654 |
| Convolution | 3×3×512 | 3 | 1 | 995 | 4×4×512 | 3 | 1 | 750 |
| Max pool | 1×1×512 | 2 | 2 | 1059 | 2×2×512 | 2 | 2 | 798 |
| Fully connected layers for classification |  |  |  |  |  |  |  |  |

**Figure 5** shows that the best predictive accuracy was obtained at a much earlier processing stage than the original version of VGG-19, as both modified VGG-19 models appeared to peak between the processing stages of conv1_1 and conv2_1, much early than was observed for the original version of VGG-19. Although these models did not attain the level of neural predictivity of VGG-19, their within-network patterns of performance provide further evidence in favor of the interpretation that only a small number of non-linear computations are needed to account for V1 neuronal responses.

**Figure 5.**
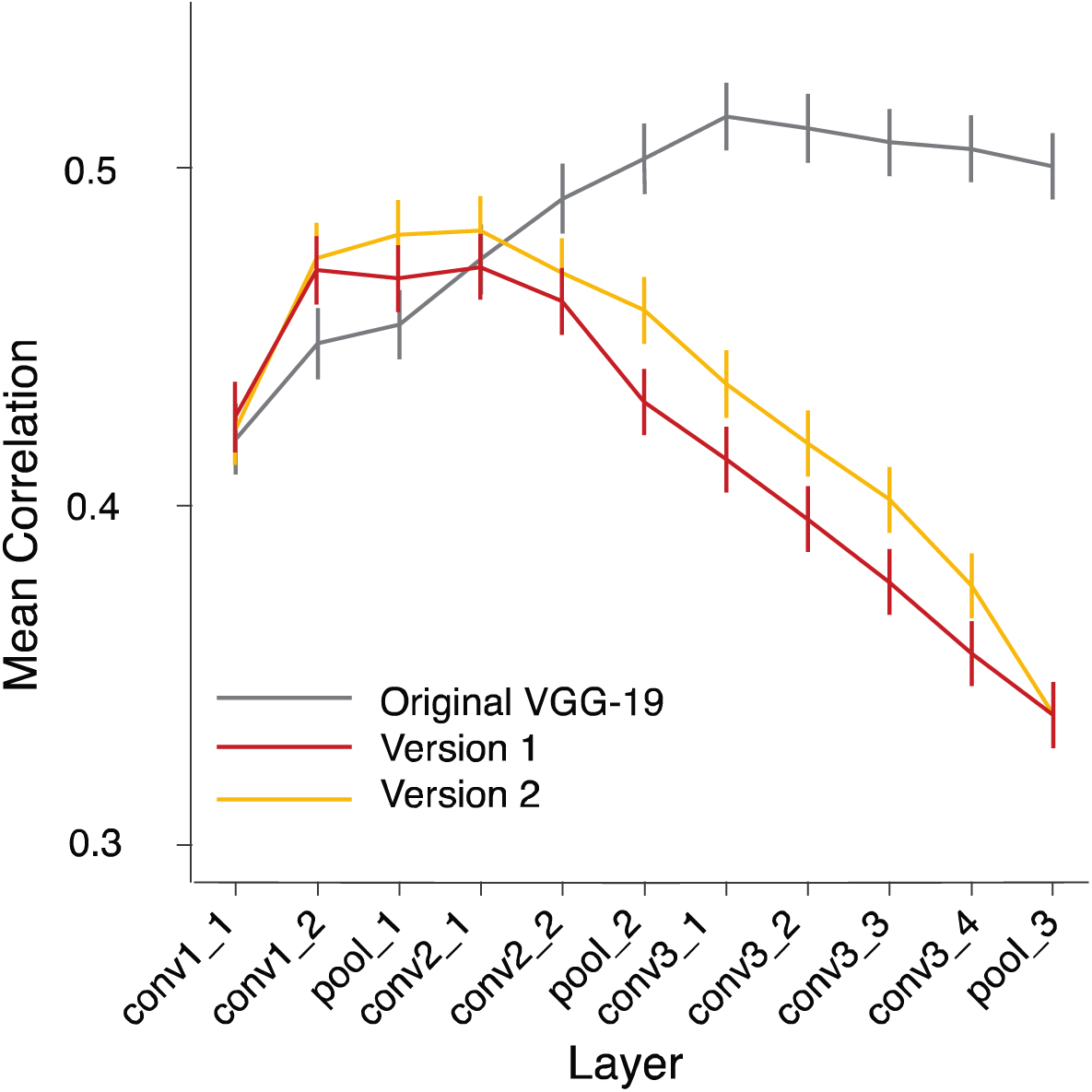
Predictive performance of the modified VGG-19 models. The mean correlation between predicted and actual V1 responses peaked in the third convolutional layer (conv2_1) for modified VGG-19 Version 1 and also Version 2.

### Evaluation of V1 Gabor pyramid models with non-linearities

Given that the best-performing layer of pre-trained AlexNet emerged in convolutional layer 1, this suggests that much simpler *shallow* models have the potential to perform well at accounting for the response properties of V1 neurons. Whereas AlexNet acquires its front-end filters from supervised training at object classification, another approach could be to use hand-engineered filters to model V1 responses, as one might be able to choose a more evenly distributed or complete set of basis filters.

We were therefore motivated to perform a parallel set of analyses on this V1 dataset using a Gabor-based pyramid model (Kay et al., 2008; Lee, 1996) to test the predictive performance of a base model and evaluate whether additional non-linear operations are needed to account for V1 responses to complex natural images. Our Gabor pyramid consisted of an array of simulated simple cell and complex cell responses that spanned four spatial scales, eight orientations, and four spatial phases (see **Methods**). This array of simulated units was used to calculate response patterns to each image, after which regularized regression was used to predict the responses of individual V1 neurons. Our evaluation of the base V1 Gabor model revealed that it predicted V1 responses quite well, and actually exhibited a marginally significant advantage when compared with conv1 of pre-trained AlexNet (mean *r* = 0.4946 vs. 0.4887, *t*(165) = 1.92, *p* = 0.057).

Moreover, it is well documented that V1 responses increase monotonically as a function of stimulus contrast but in a non-linear manner, as response saturation or compression occurs at higher stimulus contrasts (Albrecht & Geisler, 1991; Boynton et al., 1999; Ohzawa et al., 1985; Sclar et al., 1990; Skottun et al., 1991; Tong et al., 2012). Thus, one of the simplest forms of non-linearity that can be applied to a modeled V1 response, after half-wave rectification, is some type of contrast saturation effect. We mimicked the effects of contrast saturation by applying an exponentiation function to the output response of modeled simple cell and complex cell units, using exponent values ranging from 0.1 to 1.0 with increments of 0.1. This analysis revealed that compression of the Gabor-based responses with an exponent value of 0.6 (i.e., close to a square root function) led to a quantitatively modest but statistically significant improvement in the prediction of V1 responses when compared to the base model with no compression (mean *r* of 0.4979 vs. 0.4946 respectively, *t*(165) = 3.94, *p* = 1.22 ×10^-4^). Moreover, this version of the Gabor model performed significantly better than conv1 of pre-trained AlexNet (*t*(165) = 3.13, *p* = 0.002). While our Gabor model with contrast saturation did not quite reach the predictive accuracy of the best-performing layer of VGG-19 or modified AlexNet, the differences in performance between these CNN models and our Gabor model were quite modest, with CNNs showing only a modest advantage as the differences in mean *r* fell below a value of 0.02 (Figure 6).

**Figure 6.**
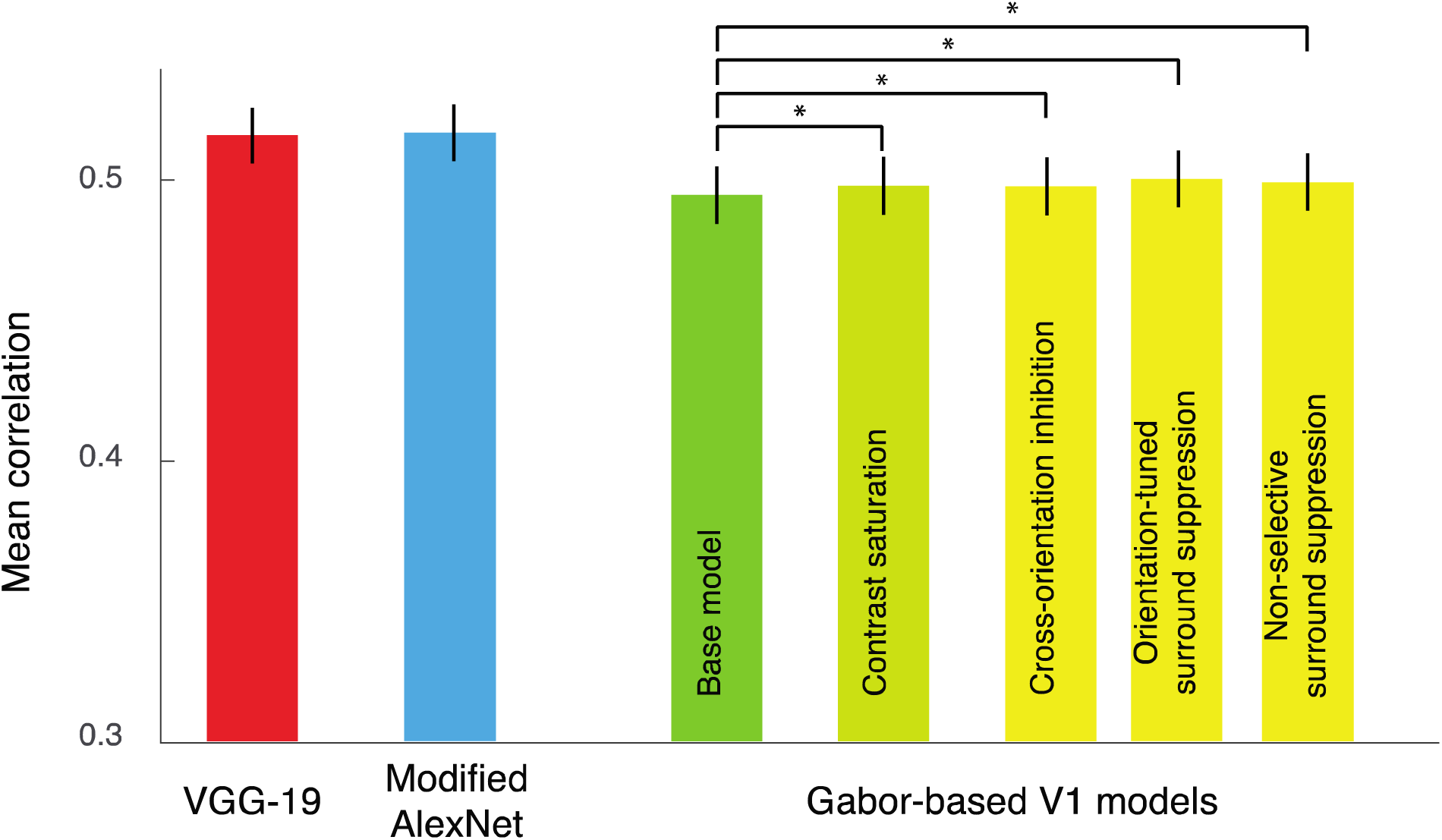
Comparison between CNN-based V1 models and Gabor-based V1 models. Bar plots show the predictive accuracy of conv3_1 layer of VGG-19, pool_1 layer of modified AlexNet, and multiple versions of the Gabor-based V1 model.

The effects of both cross-orientation inhibition and orientation-tuned surround suppression have been extensively studied in V1 (Bair et al., 2003; Bonds, 1989; Busse et al., 2009; Cavanaugh et al., 2002; Deangelis et al., 1994; Deangelis et al., 1992; Jones et al., 2001; Morrone et al., 1982; Nurminen et al., 2018; Poltoratski & Tong, 2020; Vinje & Gallant, 2002), and are commonly implemented in computational models by assuming divisive normalization (Carandini et al., 1997; Heeger, 1992b). Divisive normalization is believed to reflect a canonical neural computation that occurs in most brain areas, in which the feedforward response of each excitatory neuron is divisively modified by the summed activity of its neighbors, presumably as a consequence of some form of local inhibition (Carandini & Heeger, 2012). While cross-orientation inhibition and surround suppression can be readily demonstrated by using artificial stimuli that are specifically tailored to test for these effects (e.g., sinewave gratings), it is unclear whether such modulatory interactions would be readily detectable when evaluating V1 responses to a large diverse set of natural (and synthetic) images.

For our first set of analyses, we constructed V1 Gabor models with different types of divisive normalization. One model mimicked cross-orientation inhibition by applying normalization across all oriented units of a given spatial scale in a location-specific manner. Another model mimicked the effects of surround suppression by applying normalization across spatially neighboring units (using a 2D Gaussian window) that shared a common orientation and spatial frequency preference, while a third model applied spatial normalization without selectivity for orientation. For each model, a constant additive term in the denominator was allowed to vary as a free parameter to modify the strength of divisive normalization (see **Methods**).

In comparison to the base model (mean r = 0.4946), we found that normalization led to significantly better performance for the cross-orientation inhibition model (mean *r* = 0.4976, *t*(165) = 4.32, *p* = 2.64 ×10^-5^), the orientation-tuned surround suppression model (mean *r* = 0.5002, *t*(165) = 7.39, *p* = 7.12 ×10^-12^), as well as the non-selective surround suppression model (mean *r* = 0.4992, *t*(165) = 6.33, p = 2.21 ×10^-9^). While the above findings provide tentative evidence that normalization may affect V1 neuronal responses to complex images, it is important to consider whether normalization would further improve the prediction of V1 responses after the non-linear effect of contrast saturation is taken into consideration. We therefore implemented the contrast saturation effect with an exponent of 0.6, and then performed divisive normalization with the models described above. In comparison to the contrast saturation model without normalization (mean *r* = 0.4979), the model with both contrast saturation and cross-orientation inhibition did not show significant gains in performance (mean *r* = 0.4977, *t*(165) = 0.58, p = 0.56). Likewise, the model with contrast saturation and orientation-tuned surround suppression did not outperform the model with contrast saturation alone (mean *r* = 0.4982, *t*(165) = 0.2869, p = 0.77). However, we did observe significant improvement in V1 predictions for the contrast-saturated model with non-selective surround suppression (mean *r* = 0.4991, *t*(165) = 2.17, p = 0.03) when compared to the contrast saturation model without normalization. One interpretation of these findings is that normalization is less influential if one pre-supposes the existence of a mechanism to mediate contrast saturation in V1 neurons. However, an alternative interpretation is that some form of normalization (e.g., cross-orientation inhibition) is responsible for causing contrast saturation in V1, as has been suggested by prior research (Heeger, 1992b).

Our analyses of these V1 responses using multiple variants of a Gabor pyramid model provide positive evidence of additional non-linear computations that take place prior to or within V1, which remain detectable in neural responses to complex natural images (Coen-Cagli et al., 2015; Vinje & Gallant, 2002). It is also worth noting that while the potential contributions of divisive normalization appear quite modest in this study of V1 responses to natural and complex images, much more powerful effects have been documented in studies that rely on artificial stimuli (e.g., gratings) and tailored experimental conditions to test for the effects of cross-orientation inhibition and surround suppression (e.g., (Busse et al., 2009; Cavanaugh et al., 2002)). Thus, whether it is best to investigate the response properties of V1 using natural images or artificial stimuli may depend on the nature of the neuroscientific question to be tested (Felsen & Dan, 2005; Kay et al., 2008; Rust & Movshon, 2005).

Finally, while our best-performing Gabor model with orientation-tuned surround suppression could predict V1 responses suitably well (mean *r* = 0.5002), it was still outperformed by VGG-19 (*r* = 0.5158) and modified AlexNet (*r* = 0.5167) by a modest but statistically significant margin. Given that these V1 Gabor models are arguably much simpler, more parsimonious and more readily interpretable, how should these factors be weighted in comparison to the more complex and less readily interpretable CNN models? The preferred model of choice may depend on the goals of the study or the preferred theoretical framework of the researcher. From our own perspective, we suspect that the simpler Gabor-based model with additional normalization mechanisms may account for a majority of V1 tuning properties, while a subset of V1 neurons may have more complex receptive fields that arise from a longer sequence of non-linear operations. Whether it might be possible to adjudicate between best-fitting models at the single-neuron level remains a potentially interesting and challenging question for future studies to explore.

## Discussion

Convolutional neural networks have become increasingly influential in neuroscientific research, as they currently provide the best models for predicting neural responses to complex natural images in both lower and higher visual areas (Bankson et al., 2018; Bashivan et al., 2019; Cadena et al., 2019; Cichy et al., 2016; Guclu & van Gerven, 2015; Horikawa & Kamitani, 2017; Jang et al., 2021; Kar et al., 2019; Khaligh-Razavi & Kriegeskorte, 2014; Kietzmann et al., 2019; Nonaka et al., 2021; Schrimpf et al., 2018; Tong & Jang, 2021; Xu & Vaziri-Pashkam, 2021; Yamins & DiCarlo, 2016; Yamins et al., 2014). Understanding how these CNN models attain such high levels of neural predictivity, especially when compared to simpler models, is important for appreciating the neurally relevant computations performed by CNNs and for understanding what other factors might account for their gains in performance. Here, we investigated the claims of a recent study that reported obtaining the best predictions of V1 neuronal responses by using the unit activity patterns from an intermediate layer of VGG-19 after multiple non-linear operations were performed on the initial input (Cadena et al., 2019).

Although it is possible that stimulus-driven responses in V1 involve several non-linear operations, we hypothesized that units in the lower layers of VGG-19 may have led to poorer predictions of V1 neuronal responses because of their small receptive field size when compared to the input images used for model evaluation. We obtained multiple lines of evidence to support this view. First, we evaluated AlexNet, a well-known CNN with much larger convolutional filters in the early layers. The effective receptive field size of AlexNet rapidly increases over the first few layers, making it suitable to adjudicate our question. We found that AlexNet achieved its best predictive performance by the first convolutional layer. Although layer conv3_1 of VGG-19 performed somewhat better than this version of AlexNet, we found that a modified version of AlexNet was capable of matching the predictive performance of VGG-19, and here the best-performing layer emerged in the first pooling layer after only two prior stages of convolutional processing. These findings provide compelling evidence that while V1 is not strictly linear, the number of non-linear computations needed to explain stimulus-driven V1 responses is far less than recently claimed by Cadena et al. and in much better agreement with other prominent V1 models (Heeger, 1992a; Jones & Palmer, 1987; Rust et al., 2005; Vintch et al., 2015).

To better understand how CNN model performance may depend on the relationship between input stimulus size and the effective receptive field size of the units in a given layer used for neural prediction, we performed a set of follow-up analyses. In a control analysis performed on VGG-19, we found that smaller input images caused the best predictions to shift to lower layers of VGG-19 whereas larger input images caused a shift towards higher layers. (Although Cadena et al. (2019) performed a similar control analysis, their image size manipulations of scale 1.5 were too modest to lead to significant shifts in performance accuracy.) These findings provide compelling evidence that the relationship between CNN receptive field size and input image size can directly impact how well CNNs can predict neuronal responses. In related work, Marques et al. (2021) reported that the choice of field of view size could affect the estimated similarity between CNN unit responses and recorded V1 neuronal responses, including properties such as their size and spatial frequency tuning. Presumably, if too many units must be considered to account for the full spatial extent of a V1 neuron’s receptive field, the resulting model fits may become less stable or less accurate.

In a final set of analyses, we constructed modified versions of VGG-19 with increased stride values in the lower layers so that larger receptive fields would be acquired from supervised training on object classification. These modified versions of VGG-19 attained their highest level of V1 predictivity at an earlier processing stage than that of the original VGG-19 model (i.e., conv3_1). Thus, manipulations of CNN receptive field size and stimulus input size can both cause systematic shifts in the CNN layer that best predicts neuronal responses in early visual areas, with the best performance typically observed in a CNN layer with receptive fields that are modestly smaller than the input images used to evoke model responses.

In addition to our evaluation of CNN models, we compared different versions of a Gabor wavelet pyramid model to gain a better understanding of the non-linear tuning properties of V1 neurons. Although the basic Gabor pyramid model was not able to attain the predictive accuracy of the best-performing layer of AlexNet or VGG-19, its performance was still quite high despite the model’s simplicity. First, we confirmed that V1 neurons showed evidence of contrast saturation in their responses to the complex natural and synthetic images, consistent with previous work that tested simpler stimuli (Albrecht & Geisler, 1991; Boynton et al., 1999; Ohzawa et al., 1985; Sclar et al., 1990; Skottun et al., 1991; Tong et al., 2012). Next, we tested for potential effects of cross-orientation inhibition and surround suppression by incorporating various types of divisive normalization (Carandini & Heeger, 2012) into our Gabor pyramid model. We found clear evidence indicating that divisive normalization from neighboring local units (irrespective of orientation) led to better predictions of V1 responses. Also, if we did not presuppose a separate mechanism to mediate contrast saturation, we found that better model performance arose by incorporating mechanisms of cross-orientation inhibition and orientation-tuned surround suppression into our V1 Gabor models. These findings indicate that the effects of surround suppression in V1 are pervasive enough to be detected when evaluated using an arbitrarily selected set of complex natural images. That said, the use of artificial images such as sinewave gratings may be more effective for generating tailored stimuli to test specific hypotheses about the response properties of V1 (Rust et al., 2005), including the nature of excitatory center / suppressive surround interactions (Mely et al., 2018).

To provide a full evaluation of the complexity of neuronal tuning in V1, it is necessary to consider both simple and more complex computational models. CNNs can be very informative in this regard as they are designed to simulate the feedforward operations of the visual system. Also, CNNs learn useful statistical properties from natural images and moreover, a single network will acquire multi-level representations of varying complexity that can be used to assess neuronal complexity. However, as we show here, it is critically important to consider the impact of other correlated variables such as unit receptive field size when one uses CNNs to evaluate the complexity of neuronal tuning. We conclude that at least during the first wave of feedforward processing, V1 neuronal responses are well described by simple linear filters followed by only a few non-linear operations.

Our findings are consistent with traditional V1 models that have typically relied on only 1-2 non-linear operations to fit V1 neuronal responses (Jones & Palmer, 1987; Rust et al., 2005; Vintch et al., 2015). Why then do CNN models tend to outperform these traditional V1 models, including the V1 Gabor pyramid model that was tested here? One possibility might be that CNNs learn better filters in layer 1 due to their training with large sets of natural images. However, our base V1 Gabor pyramid model performed significantly better than layer 1 of pre-trained AlexNet. Instead, the gains in CNN performance appeared to occur after layer 1, after which the units can acquire more complex receptive field structures by combining the responses from multiple initial filters with varying shape, orientation and spatial frequency tuning. However, given the modest margin by which the CNN models outperformed our Gabor-based models, it is conceivable that only a fraction of the neurons in V1 acquire these more complex non-linear receptive fields while the majority of V1 neurons can be well described by a weighted set of inputs from a Gabor-based pyramid model.

Finally, it should be emphasized that Cadena et al.’s study was primarily focused on characterizing the feedforward or stimulus-driven aspects of V1 neuronal responses. Other studies have shown that recurrent visual processing and top-down effects of task-based attention can have powerful modulatory effects on neuronal responses in macaque V1 (Gilbert & Li, 2013; Lamme & Roelfsema, 2000; Roelfsema et al., 1998). Work from our own research group has shown that binocular rivalry competition, spatial and feature-based attention, object-based attentional selection, and visual working memory, all lead to powerful modulatory effects in the human primary visual cortex (Cohen & Tong, 2015; Harrison & Tong, 2009; Jehee et al., 2011; Kamitani & Tong, 2005; Tong & Engel, 2001). It will be of considerable interest for future studies to explore whether variations in CNN architecture, the incorporation of recurrent or top-down processing or the expansion of stimuli and methods used for network training can further improve the ability of CNN models to predict the non-linear response properties of V1.

## Acknowledgments

This work was supported by the National Institutes of Health, National Eye Institute Awards R01EY029278 and R01EY035157 to FT, and a P30EY008126 center grant to the Vanderbilt Vision Research Center. The authors thank Hojin Jang for input on the CNN analysis and Dave Coggan for feedback on early versions of this manuscript.

